# The circulating phageome reflects bacterial infections

**DOI:** 10.1101/2022.08.15.504009

**Authors:** NL Haddock, LJ Barkal, N Ram-Mohan, G Kaber, CY Chiu, AS Bhatt, S Yang, PL Bollyky

## Abstract

Bacteriophage, viruses that infect bacteria, are abundant in the human body but the relationship between the phageome and bacterial population dynamics is unclear. Because bacteriophage are often highly specific to bacterial host strains and species, we asked whether bacteriophage present in cell-free DNA (cfDNA) reflect bacterial infections in sepsis. To address this, we generated a workflow for identifying and interpreting bacteriophage sequences in cfDNA and a bacteriophage characteristic dictionary. In two independent cohorts of infected patients and asymptomatic controls, we demonstrate that all individuals, septic and healthy, have a circulating phageome. Moreover, infection associates with overrepresentation of pathogen-specific phage, allowing for the study of bacterial pathogens. We further show that phage can identify pathovariant *Escherichia coli* infections and distinguish between closely-related pathogenic bacterial species such as *Staphylococcus aureus* and frequent contaminants such as coagulase-negative Staphylococcus. Phage DNA may have utility in studying bacteriophage ecology in infection.

## Introduction

Bacterial culture methods developed over a century ago remain the standard of care for identifying bacterial pathogens in sepsis and other settings. Unfortunately, these methods are often imprecise and can be confounded by both false positives and negatives. Improving the identification and subsequent study of bacterial pathogens is a critical medical need^1^.

Next generation sequencing (NGS) of cell-free DNA (cfDNA) holds promise for filling this gap. cfDNA is made up of DNA fragments, typically 50-200 bp long, found in circulation^2^. cfDNA is primarily human in origin with a small but rich microbial compartment^2–4^. With high-throughput sequencing, cfDNA lends itself to non-invasive testing and has transformed diagnostics in perinatal testing, autoimmune disorders, cancer staging, and transplant rejection^5–8^. There have been efforts to develop cfDNA infection diagnostics in sepsis^9–12^ and other settings^13, 14^.

Retrospective cohort studies show that these approaches failed to identify the pathogen in many sepsis cases, and these approaches do not establish thresholds distinguishing colonization from infection^15, 16^. Other molecular approaches such as PCR-based diagnostics yield similarly mixed results^17^ and have limited widespread adoption. There is an urgent need for better methods to study bacterial infection ecology and facilitate downstream diagnostic improvements.

Bacteriophage (phage), or viruses that infect bacteria, may provide insight into the bacterial ecology underlying sepsis and opportunistic infections. Phage are present in all compartments of the human body, including the skin and gut. Here they infect bacterial populations and are an active and critical component of the human microbiome^18, 19^. Though abundant in the human body, they have historically been challenging to identify in samples and have been termed microbial “dark matter”^20, 21^.

Nonetheless, aspects of phage biology make them uniquely well-suited for interrogating bacterial infections. Many phage have narrow host ranges limited to a few closely related bacteria, though information is sparse given the vast diversity of phage^22–24^. Composition of phages may therefore reflect bacterial populations at the species and strain level. Additionally, some phage translocate across epithelia and into the bloodstream^18, 25, 26^. Despite structural and genetic diversity, most phage within humans are non-enveloped DNA viruses^27, 28^ and any phage DNA entering circulation can be sequenced using existing NGS methods and detected in cfDNA^3, 29^.

Targeted analysis of phage in metagenomic data has historically been limited by lack of available phage sequences^20, 30–32^. Additionally, many metagenomic annotation software packages require upstream metagenome assembly that is frequently impossible in low microbial biomass specimens. Fortunately, continuous rapid increases in submitted genomes to NCBI’s GenBank has facilitated improved identification of these phage as more of these “dark matter” genomes become known. Additionally, there are ongoing efforts to better classify phage taxonomically via genomic methods rather than the tradition of grouping only by phage particle morphology^33^.

Here we describe an approach to confidently identify and interpret DNA phage sequences in cfDNA. We apply this workflow to plasma samples collected from two independent cohorts, each comprised of patients with infection-driven sepsis as well as healthy controls. One of these cohorts involves previously published metagenomic and culture data^34^ while the other was generated as part of this work. We reveal that phageomes reflect infection through pathogen-specific overrepresentation.

Furthermore, we show that phage can distinguish between closely related bacterial species where bacterial cfDNA may fall short. Our data suggest that circulating phage sequences in plasma provide a non-invasive approach to studying bacteriophage ecology in infection.

## Results

### The circulating phageome is mostly unaltered by sepsis but reflects infection etiology

We sought to determine if there is a circulating phageome in healthy subjects and whether it is disrupted in patients with blood culture positive infections. To this end, we analyzed plasma cfDNA from 61 patients with sepsis seen in the Stanford University Emergency Department as well as 10 asymptomatic controls (Fig. 1A). The sepsis samples represent twenty different bacterial infections confirmed by positive blood cultures, and three of the samples have positive cultures for multiple bacterial genera (Table 1). The samples were analyzed using the workflow depicted in Fig. 1A. Briefly, cfDNA was collected from the samples and sequenced on an Illumina platform to an average depth of 18.67 million reads. Raw data was quality controlled and trimmed using standard bioinformatics tools, FASTQC^35^ and Trimmomatic^36^. Human reads were subtracted by mapping to the human reference genome GRCh38 via Bowtie2.

**Figure 1.**
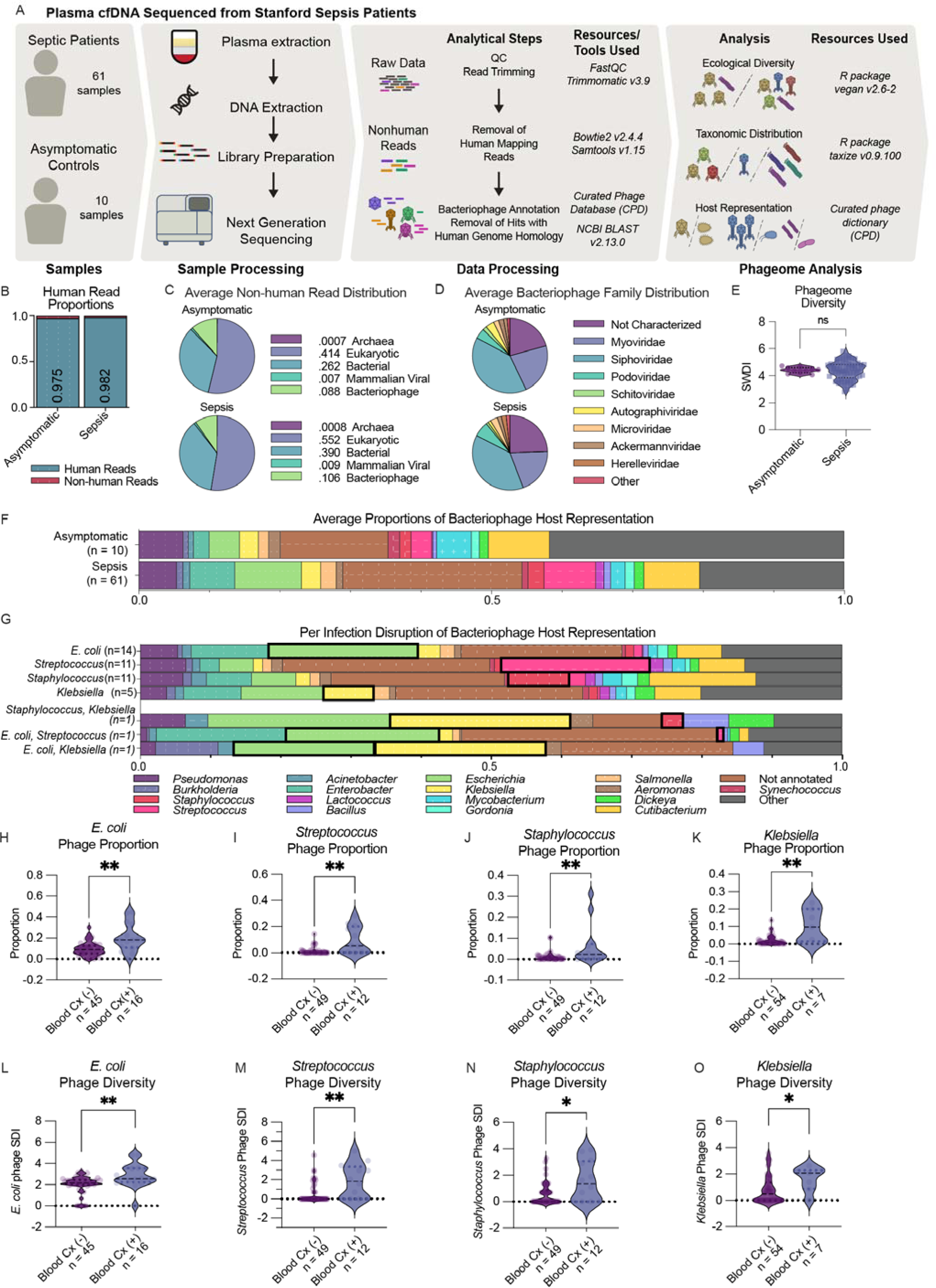
The circulating phageome is mostly unaltered by sepsis but reflects infection etiology. A) Schematic of sample numbers, sample processing, and analytical approach. B) Proportions of human and non-human reads. C) Distribution of non-human reads identity by Archeal, Eukaryotic, Bacterial, Mammalian Viral, and Bacteriophage categories. Asymptomatic (Archaea mean 0.0007 SD 0.0003, Eukaryotic mean 0.420 SD 0.126, Bacterial mean 0.486 SD 0.126, Mammalian Viral mean 0.006 SD 0.002, Bacteriophage mean 0.0877 SD 0.0223) Sepsis (Archaea mean 0.0008 SD 0.0005, Eukaryotic mean 0.552 SD 0.275, Bacterial mean 0.390 SD 0.281, Mammalian Viral mean 0.009 SD 0.018, Bacteriophage mean 0.1065 SD 0.083) D) Distribution of bacteriophage distribution of bacteriophage family. E) Shannon Diversity Index in asymptomatic and septic patient phageomes is not significantly different (Asymptomatic: mean = 4.41, range = 0.0655, Sepsis: mean = 4.38, range = 2.992, P = 0.943 per Mann-Whitney test) shown by violin plot with median and quartiles shown by dashed lines. F) Average distribution of unique phages by bacterial host genus in asymptomatic and septic patient phageomes. In Asymptomatic samples, mean proportions and SD (Pseudomonas: mean 0.059, SD 0.035, Acinetobacter: mean 0.007 SD 0.004, Escherichia: mean 0.043 SD 0.021, Salmonella: mean 0.015 SD 0.013, Not Annotated: mean 0.16 SD 0.06, Burkholderia: mean 0.0063 SD 0.0060, Enterobacter: mean 0.0024 SD 0.034, Klebsiella: mean 0.024 SD 0.027, Aeromonas: mean 0.017 SD 0.011, Synechococcus: mean 0.013 SD 0.033, Staphylococcus: mean 0.015 SD 0.027, Lactococcus: mean 0.0019 SD 0.0045, Gordonia: mean 0.012, SD 0.016, Cutibacterium: mean 0.088 SD 0.083, Streptococcus: mean 0.024 SD 0.037, Bacillus: mean 0.057 SD 0.053, Mycobacterium: mean 0.048 SD 0.047, Dickeya: mean 0.013, Other Host: mean 0.42 SD 0.065). In Sepsis samples, mean proportions and SD (Pseudomonas: mean 0.0582 SD 0.073, Acinetobacter: mean 0.017 SD 0.024, Escherichia: mean 0.095 SD 0.080, Salmonella: mean 0.015 SD 0.018, Not Annotated: mean 0.23 SD 0.11, Burkholderia: mean 0.0093 SD 0.012, Enterobacter: mean 0.052 SD 0.062, Klebsiella: mean 0.030 SD 0.046, Aeromonas: mean 0.014 SD 0.014, Synechococcus: mean 0.0069 SD 0.016, Staphylococcus: mean 0.024 SD 0.054, Lactococcus: 0.015 SD 0.054, Gordonia: mean 0.012 SD 0.021, Cutibacterium: mean 0.058 SD 0.086, Streptococcus: mean 0.031 SD 0.064, Bacillus: mean 0.016 SD 0.018, Mycobacterium: mean 0.021 SD 0.040, Dickeya: 0.018 SD 0.014, Other Host: mean 0.28 SD 0.14). G) Average proportions of unique phage by bacterial host in samples with single identified pathogen, by cultured pathogen (Streptococcus, Staphylococcus, Pseudomonas, Klebsiella, and Escherichia) as well as three polymicrobial infections with bar corresponding to row’s infectious pathogen outlined in black outline. H-K) Infection specific phage proportion shown as violin plots with median and quartiles shown by dashed lines. Summary Statistics of Mann-Whitney tests available in Fig.S5 A. H) E. coli infection status (P = 0.0017). I) Streptococcus infection status (P = 0.0018). J) Staphylococcus infection status (P = 0.0094). K) Klebsiella infection status (P = 0.0056). L-O) Infection specific Shannon DI calculated and shown as violin plots with median and quartiles shown by dashed lines. Summary Statistics of Mann-Whitney tests available in Fig. S5B. L) E. coli infection status (P = 0.0051). M) Streptococcus infection status (P = 0.0025). N) Staphylococcus infection status (P = 0.0192). O) Klebsiella infection status (P = 0.0272). All statistics were performed using unpaired, two tailed Mann-Whitney tests.

**Table 1:** Distribution of blood culture positive bacterial identifications in the sequenced sepsis cohort (N = 61).

|  |  |
| --- | --- |
| *Cronobacter sakazakii* | 1 |
| *Enterobacter aerogenes* | 2 |
| *Enterobacter cloacae* | 1 |
| *Enterococcus* sp. | 1 |
| *Escherichia coli* | 14 |
| *Escherichia coli* and *Klebsiella pneumoniae* | 1 |
| *Escherichia coli* and *Streptococcus* sp. | 1 |
| Gram Positive cocci | 1 |
| *Klebsiella pneumoniae* | 5 |
| *Moraxella* sp. | 1 |
| *Paenibacillus* sp. | 1 |
| *Proteus vulgaris* | 1 |
| *Pseudomonas aeruginosa* | 6 |
| *Salmonella enterica* | 1 |
| *Serratia marcescens* | 1 |
| *Staphylococcus aureus* | 5 |
| *Staphylococcus aureus* and coagulate-negative *Staphylococcus* sp. | 1 |
| *Staphylococcus aureus* and *Klebsiella pneumoniae* | 1 |
| *Staphylococcus* sp., coagulase-negative | 5 |
| *Streptococcus* sp. | 11 |
| *Streptococcus pneumoniae* | (2) |
| *Streptococcus mitis* | (2) |
| *Streptococcus pyogenes* | (1) |

Plasma cfDNA from both asymptomatic controls and septic patients was comprised of mostly human reads (Fig. 1B), consistent with previous studies^2–4^. A BLAST search utilizing the full NCBI Nucleotide database revealed that non-human reads, despite making up a relatively small proportion of all cfDNA reads, represented a rich microbial community (Fig. 1C). Bacterial hits include many genera. In asymptomatic individuals the greatest single genus represented was *Cutibacterium*, a known skin commensal^37^ (Fig. S1A). Less than 1% of reads corresponded to mammalian viruses, while considerably more reads mapped to bacteriophage: 8.80% and 9.62% of non-human reads in asymptomatic and septic patient samples, respectively (Fig. 1C, Fig S1B), and all samples contained phage reads.

A first-pass BLAST search^38, 39^ was performed against our Curated Phage Database (CPD), which was constructed from the NCBI Nucleotide database. Reads with significant hits were subjected to a secondary and more stringent human sequence removal, in which all reads with significant BLAST hits to any human sequence deposited in the NCBI nuccore were removed and annotations were made again using the CPD.

A common challenge in phage research is linking phage with the known bacterial host(s). The CPD pulls from virushostDB^40^, naming conventions of bacteriophages with clearly identified host genera, and NCBI nucleotide source host fields for bacteriophage sequence entries to include available bacterial host metadata. Taxonomic classifications for both phage and their bacterial host(s) are also included as metadata in the CPD, which we have made publicly available (https://doi.org/10.5281/zenodo.7154236). Additional human depletion steps and the CPD allow for clean phageome annotations that can be meaningfully analyzed with respect to infection etiology.

Using more stringent human read depletion, fewer human reads interfered with phage calls (Fig. S2A) and annotations had fewer phage without known bacterial hosts (Fig. S2B). Remaining annotations reflect a broad variety of phage families (Fig. 1D), though the tailed phage families Myoviridae and Siphoviridae predominated. There was no statistical difference in phage family representation within the circulating phageomes of asymptomatic controls and septic patients (Fig. 1D). The Shannon Diversity Index (SDI), a common ecological quantification of species richness and evenness^41^ of circulating phage was not significantly different between the groups (Fig. 1E), though septic patients did have a wider range in phage SDI. These findings indicate that there is no broad “Sepsis Phageome” signature.

Of identified phage with known bacterial hosts in the CPD, most corresponded to *Pseudomonas*, *Escherichia, Klebsiella, Cutibacterium*, and *Streptococcus* genera (Fig. 1F). Bacterial host proportions were not significantly different by sepsis status (Fig. 1F, Fig. S3A, Table S1). There was inter-individual variation with most samples dissimilar from all others on a per-phage basis and the majority of identified phage present in less than 10% of samples (Fig. S3B-F). Many phage have unknown bacterial hosts, accounting for 47.2% and 37.4% of phage in the asymptomatic and septic patient groups, respectively (Fig. 1F). A portion of these phage were uncharacterized gut phages from gut metagenome sequencing, suggesting potential contribution of gut phages to the circulating cfDNA pool. Furthermore, uncharacterized phages in asymptomatic individuals were comprised of higher proportions of these gut phages than in individuals with sepsis (Fig. S4A).

As sepsis is commonly caused by a single bacterial strain^42^, we investigated circulating phage on a per-infection basis – hypothesizing that bacteriophage pertinent to the causative pathogen in a sample will be overrepresented. We found that patients with sepsis due to *E. coli, Streptococcus spp.*, *Staphylococcus spp.*, and *Klebsiella spp.* infections had a corresponding overrepresentation in phage proportion corresponding to the causative pathogen (Fig. 1G-K, Fig. S5A). However, analysis of proportion alone does not account for overall number of phage per sample or the composition of the phage population associated with those pathogens across all samples. We therefore calculated the ecological diversity of phage. Applying SDI to each infection etiology in our dataset with at least 7 samples (a sufficient number for us to detect a 75% SDI increase with 80% power), we found increases in the SDI of the pathogen specific phage population in patients with sepsis due to *E. coli, Streptococcus spp.*, *Staphylococcus spp.*, and *Klebsiella spp.* infections (Fig. 1L-O, Fig. S5B). There were three samples with positive cultures from multiple bacterial genera that demonstrated varying degrees of representation of both pathogens (Fig. 1G, Fig. S5C).

Taken together, the content of non-human cfDNA, including the bacterial, eukaryotic, and viral DNA as well as the bacteriophage families, diversity, and annotated bacterial hosts, were broadly similar in health and in sepsis. However, septic patients with positive blood cultures had phageomes more representative of the bacterial cause of their sepsis.

### The circulating phageome reflects individual pathogens in another large study

We sought to validate our findings in a larger, more clinically diverse group of patients. The SepSeq study, a cohort study of 391 individuals, investigated the use of bacterial cfDNA to diagnose the cause of sepsis in patients triggering an emergency department sepsis alert with a wide array of infectious etiologies for which data are publicly available^34^.

There are four patient populations in the SepSeq study: patients with 1) blood culture positive sepsis 2) blood culture negative sepsis with documented infection elsewhere, 3) Systemic Inflammatory Response Syndrome (SIRS) who clinically appear septic but ultimately no infectious etiology was identified, and 4) asymptomatic controls (Fig. 2A). The original study reported overall promising diagnostic statistics using bacterial cfDNA, but with notable gaps: under-sensitivity in samples with negative initial blood cultures, and oversensitivity to chronic infections and commensals. We sought to determine first, if the phageome trends we observed in our dataset were consistent in the SepSeq study in patients with bacterial infections and second, if phage present in cfDNA might offer additional useful information in the study of pathogenic bacterial ecology.

**Figure 2.**
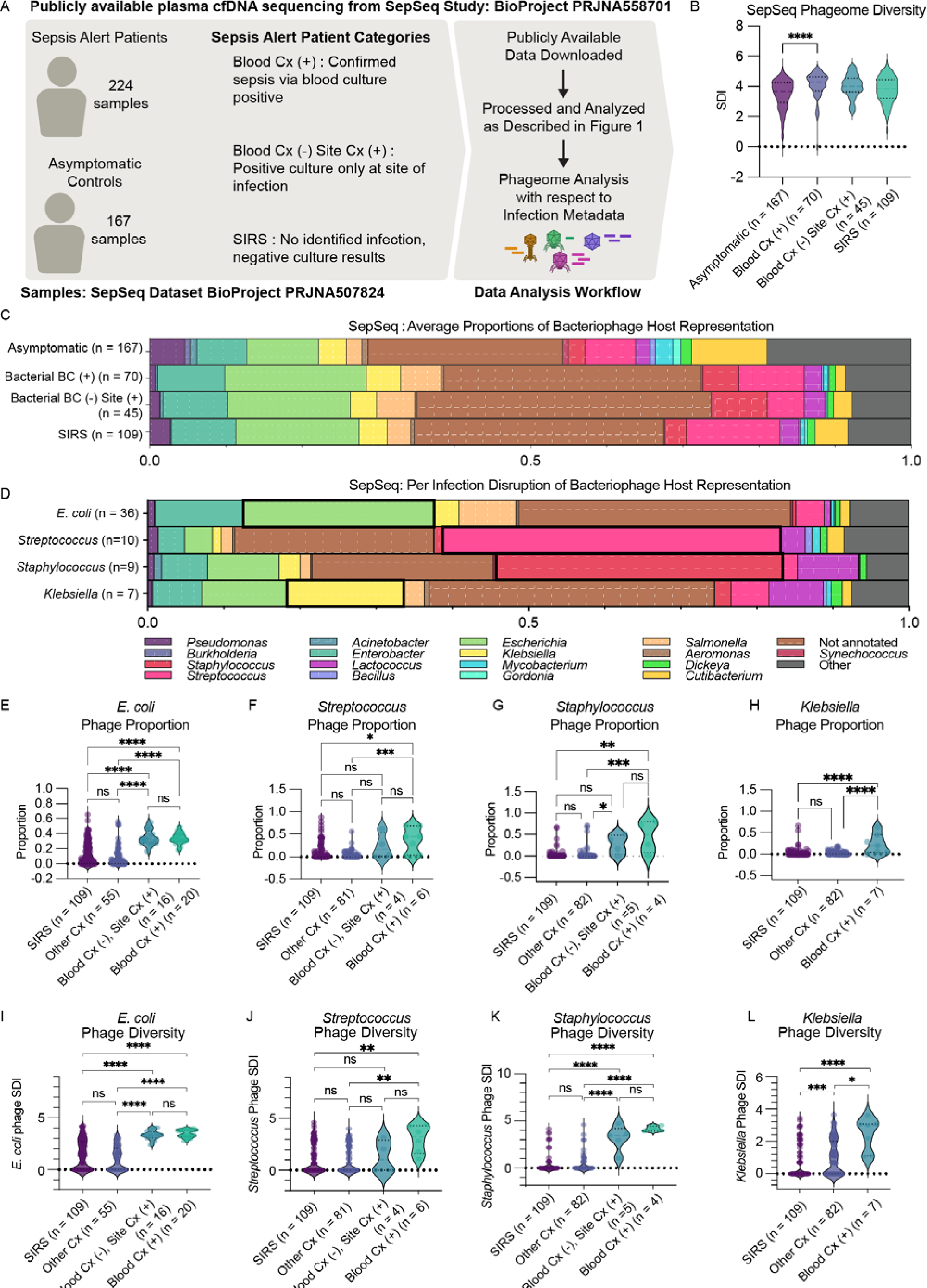
The circulating phageome reflects individual pathogens in another large study. A) SepSeq Sample Schematic. B) Shannon DI of Phages by sample category, (mean Blood Cx (+): 4.05, mean Asymptomatic: 3.48, P < 0.001). C) Average proportions of Unique Phage by bacterial host by sample category. D) Per Infection disruption of average proportions unique phage with single identified pathogen, by cultured pathogen in Streptococcus, Staphylococcus, Pseudomonas, Klebsiella, and Escherichia infection, with bar corresponding to row’s infectious pathogen outlined in black outline. (E-H) Infection specific phage proportion by infection category, all statistics performed by Kruskal-Wallis test with Dunn’s multiple comparisons shown as violin plots with median and quartiles shown by dashed lines. Summary Statistics available in Fig. S5C E) E. coli (mean SIRS: 0.152, mean Other Blood Cx: 0.096, mean Blood Cx (-) Site Cx (+): 0.347, mean Blood Cx (+): 0.338, Kruskal-Wallis P<0.0001), F) Streptococcus (mean SIRS: 0.119, mean Other Blood Cx: 0.037, mean Blood Cx (-) Site Cx (+): 0.236, mean Blood Cx (+): 0.375, Kruskal-Wallis P<0.0001), G) Staphylococcus (mean SIRS: 0.032, mean Other Blood Cx: 0.034, mean Blood Cx (-) Site Cx (+): 0.268, mean Blood Cx (+): 0.456, Kruskal-Wallis P=0.0001), H) Klebsiella (mean SIRS: 0.034, mean Other Blood Cx: 0.031, mean Blood Cx (+): 0.241, Kruskal-Wallis P=0.0002). (I-L) Pathogen Host Phage Shannon Diversity by infection category, all statistics performed by Kruskal-Wallis test with Dunn’s multiple comparisons shown as violin plots with median and quartiles shown by dashed lines. Summary Statistics available in Fig. S5D I) E. coli (mean SIRS: 1.45, mean Other Blood Cx: 1.07, mean Blood Cx (-) Site Cx (+): 3.35, mean Blood Cx (+): 3.48, Kruskal-Wallis P<0.0001), J) Streptococcus (mean SIRS: 1.02, mean Other Blood Cx: 0.760, mean Blood Cx (-) Site Cx (+): 1.32, mean Blood Cx (+): 3.03, Kruskal-Wallis P=0.0024), K) Staphylococcus (mean SIRS: 0.266, mean Other Blood Cx: 0.316, mean Blood Cx (-) Site Cx (+): 3.17, mean Blood Cx (+): 4.16, Kruskal-Wallis P<0.0001), L) Klebsiella (mean SIRS: 0.510, mean Other Blood Cx: 1.00, mean Blood Cx (+): 2.43, Kruskal-Wallis P<0.0001)

Using our annotation workflow (Fig. 2A), we again found circulating phageomes were remarkably similar in health and sepsis. While overall phage diversity was higher in patients with sepsis and positive blood cultures (Fig. 2B), proportions of bacterial hosts represented by the identified phage (Fig. 2C) were largely stable. We found the most common bacterial hosts to be *Enterobacter spp.*, *Escherichia spp.*, *Staphylococcus spp.*, and *Streptococcus spp.*, even in healthy patients (Fig. 2C) and again found that uncharacterized gut phages contribute to the phage population in these patients (Fig. S4B). Phage specific to the causal bacterial pathogen were again overrepresented in those phageomes compared to other groups (Fig. 2D, Fig. S5D). We found that in patients with blood culture positive *E. coli, Streptococcus spp.*, *Staphylococcus spp.*, and *Klebsiella spp.* infections, both the proportion (Fig. 2E-H, Fig. S5D) and diversity (Fig. 2I-L, Fig. S5E) of pathogen-associated phage were significantly higher than in individuals with another infection or no isolated pathogen (SIRS).

Interestingly, for a single bacterial infection etiology, phage proportion and diversity across patients did not vary between blood culture positive and negative groups, which indicates that circulating phage sequences are capable of reflecting infection outside the context of bacteremia. One other important point is the non-zero background of phage in individuals with SIRS or other infections – especially in the case of *Klebsiella* phage, where individuals with non-*Klebsiella* sepsis have a higher diversity of *Klebsiella* phage than in SIRS patients (Fig. 2L). These findings may be indicative of potentially undiagnosed infections, of influence of infection on other bacterial communities in the body, or even of increased phage translocation due to inflammation.

The SepSeq study redemonstrated a healthy circulating phageome and no uniform “Sepsis Phageome”. The proportion and diversity of infection-associated phage, however, was consistently overrepresented in the context of bacterial infection. This suggests plasma cfDNA sequencing provides an alternative to site of infection fluid sequencing towards study of bacterial infection ecology.

### E. coli phage reflect host strain characteristics

*E. coli* infections are common causes of sepsis, including in our cohorts, and *E. coli* phage are extensively studied. Here, *E. coli* phage were sporadic in asymptomatic patients, more common in patients with SIRS or non-*E. coli* sepsis, and ubiquitous in patients with *E. coli* sepsis (Fig. 3A). There was notable enrichment for *E. coli* phage from Siphoviridae, Podoviridae, Myoviridae phage families in *E. coli* sepsis while *Autographiviridae* phage remain at stably low levels despite infection status (Fig. 3B-E). There was also slight but significant enrichment for *E. coli* phage from Siphoviridae, Podoviridae, Myoviridae phage families in patients with SIRS or non-*E. coli* sepsis (Fig. 3B-E). Genome-based viral taxonomies are gaining traction, and testing them using a new but incomplete database^43^ results in many of the these phage being reclassified or flagged as unclassified (Fig. S6A). Still, the trends observed with the old taxonomy persist (Fig. S6B).

**Figure 3:**
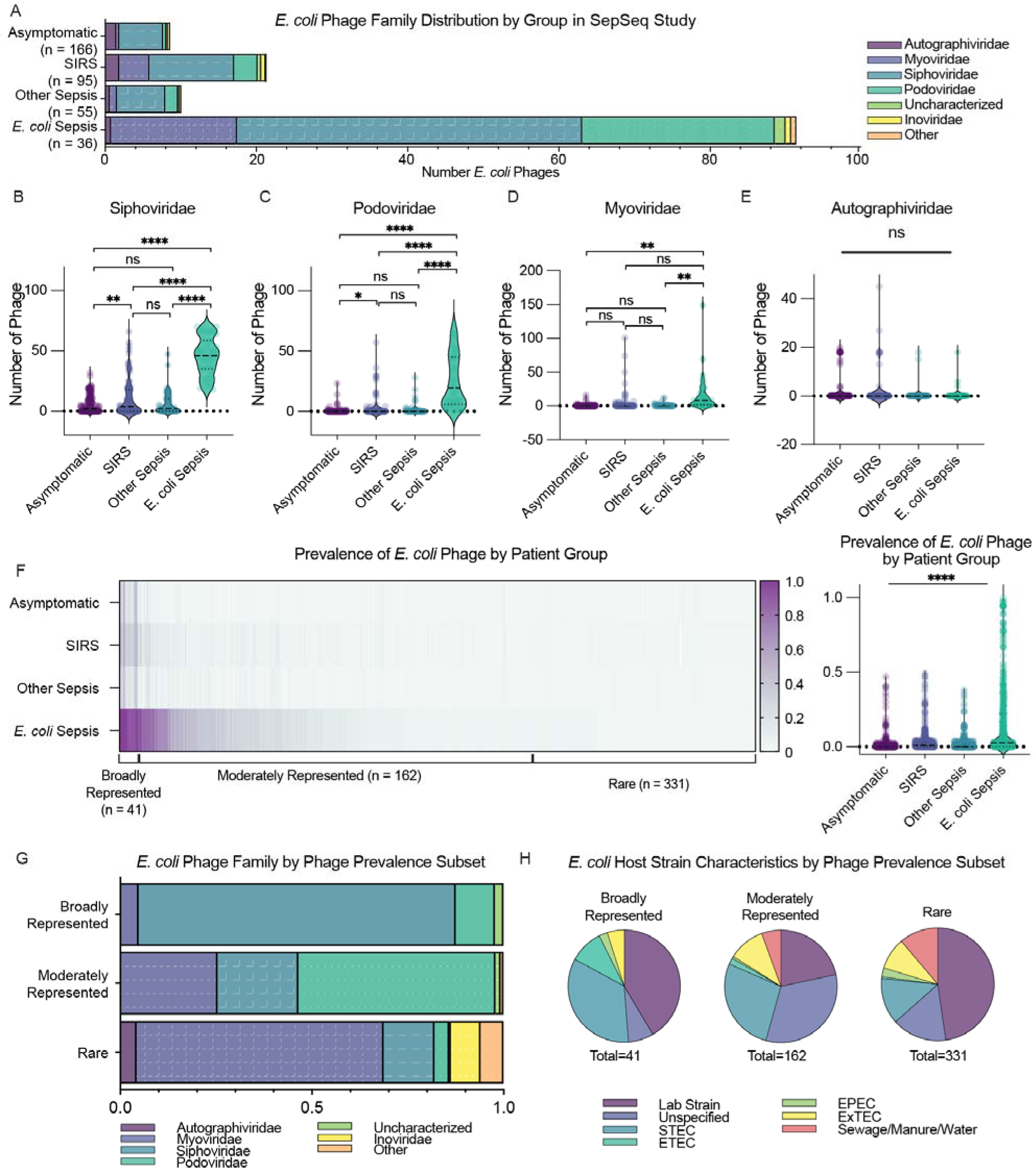
E. coli phage reflect host strain characteristics. A) In samples with E. coli phage detectable, E. coli bacteriophage numbers by family vary by infection status. (B-E) Number of E. coli phages by phage families B) E. coli Siphoviridae phages (mean Asymptomatic: 5.80, mean SIRS: 11.19, mean Other Sepsis: 6.40, mean E. coli Sepsis: 45.67, Brown-Forsythe ANOVA P < 0.0001), C) E. coli Podovidirae phages (mean Asymptomatic: 0.44, mean SIRS: 3.08, mean Other Sepsis: 1.66, mean E. coli Sepsis:, Brown-Forsythe ANOVA P < 0.0001), D) Number of E. coli Myoviridae phages (mean Asymptomatic: 0.36, mean SIRS: 4.02, mean Other Sepsis: 0.98, mean E. coli Sepsis: 16.67, Brown-Forsythe ANOVA P < 0.0001), E) E. coli Autographiviridae phages (Brown-Forsythe ANOVA test P = 0.32), F) Some individual E. coli phages are prevalent across multiple samples, with E. coli Sepsis samples reflecting phages more highly represented across the group (mean 0.1524 representation across E. coli Sepsis E. coli phages versus 0.01231 for Asymptomatic, 0.03364 for SIRS, and 0.01631 for Other Sepsis samples). G) Stacked bar plot of phage representation across samples by group of E. coli phages subset by representation level in E. coli Sepsis. H) E. coli phage morphology by phage representation groups. All violin plots have median and quartiles shown by dashed lines

There was incredible diversity of *E. coli* phage hits with more than 500 unique phage identified across the 36 *E. coli* sepsis patients. We separated these phage into three groups based on each phage’s prevalence across samples. Most strikingly, there was a group of broadly represented, or ubiquitous, *E. coli* phage present in nearly all patients with *E. coli* sepsis but not other groups. A subset of moderately represented phage were present in many but not all patients with *E. coli* sepsis. The remaining phage were present in only a few samples – in fact the majority of phage were present in only one or two patients (Fig. 3F). We then analyzed the characteristics of the individual phage by the aforementioned prevalence-based groups. We found differences in the *E. coli* phage families (and therefore morphology) that corresponded to how common the phage was amongst patients. Broadly represented *E. coli* phage possess clear enrichment for Siphoviridae phage; Podoviridae phage predominate in the moderately represented subset of phage; and the rare phage subset was predominantly Myoviridae phage (Fig. 3G).

*E. coli* phage are well studied and 65.56% of these phage have a clear bacterial host strain noted in the literature (Supplemental File 2 (Coliphage Characteristics)). Using this rich literature captured in the CPD, we annotated bacterial host to the level of the bacterial strain for the *E. coli* phage in each prevalence subset. We found that lab strains of *E. coli*, most derived initially from patient samples, were common hosts across each phage subset. Phage associated with toxin-producing strains of *E. coli* (STEC, ETEC) were overrepresented in the broadly represented group of phage (Fig. 3H) – most patients with *E. coli* sepsis had hits to these phage. A more global view of all *E. coli* phage in *E. coli* sepsis patients showed an overrepresentation of STEC associated phage (Fig. S7) with a corresponding decrease in environment-associated bacterial strains (sewage, manure, or water).

Together, these data indicate that *E. coli* phage provide an ecological view of *E. coli* infections in the context of sepsis – with phage from samples of true infection being associated with toxin producing *E. coli* host pathovariants. This level of granularity may have utility in distinguishing true infection from colonization.

### Phage can distinguish between related species Staphylococcus aureus and coagulase-negative Staphylococcus

One difficulty in plasma based microbial analyses is the fact that genomes of related bacterial species may be difficult to distinguish due to short cfDNA fragments and comparatively large genomes that may share many conserved genes. In the case of *Staphylococcus* infections, bacterial cfDNA can potentially underperform in distinguishing species within a bacterial genus. We asked the question of whether phage can help regain some of the lost “resolution” of bacterial DNA. We assessed whether bacteriophage ecology differed between the species *S. aureus*, the presence of which in blood culture is always considered pathogenic, and coagulase-negative *Staphylococcus* species (CoNS), the presence of which is often a contaminant. Both species share many core genes^44^, which if detected in cfDNA are difficult to attribute to either confidently – though culture techniques adequately distinguish these species in clinical contexts.

We found that when mapping to a *S. aureus* reference genome, bacterial cfDNA was unable to distinguish between *S. aureus* and CoNS in either our cohort (Fig. 4A) or the SepSeq cohort (Fig. 4B). Consistent with this, the corresponding ROC plots have AUCs near 0.5 (Fig. 4C). Notably, however, the populations of phage that infect *S. aureus* and CoNS are distinct and the diversity of *S. aureus*-specific phage was informative as there was a significant increase in *S. aureus* phage diversity only in *S. aureus* infections (Fig. 4E). This is reflected in a ROC plot with AUC of 0.96 (Fig. 4F). While the AUC based on phage diversity was higher than that based on bacterial DNA (Fig. 4F), this increase did not reach significance in our smaller cohort (Fig. 4D) and calls for further validation in a separate, more highly powered cohort.

**Figure 4.**
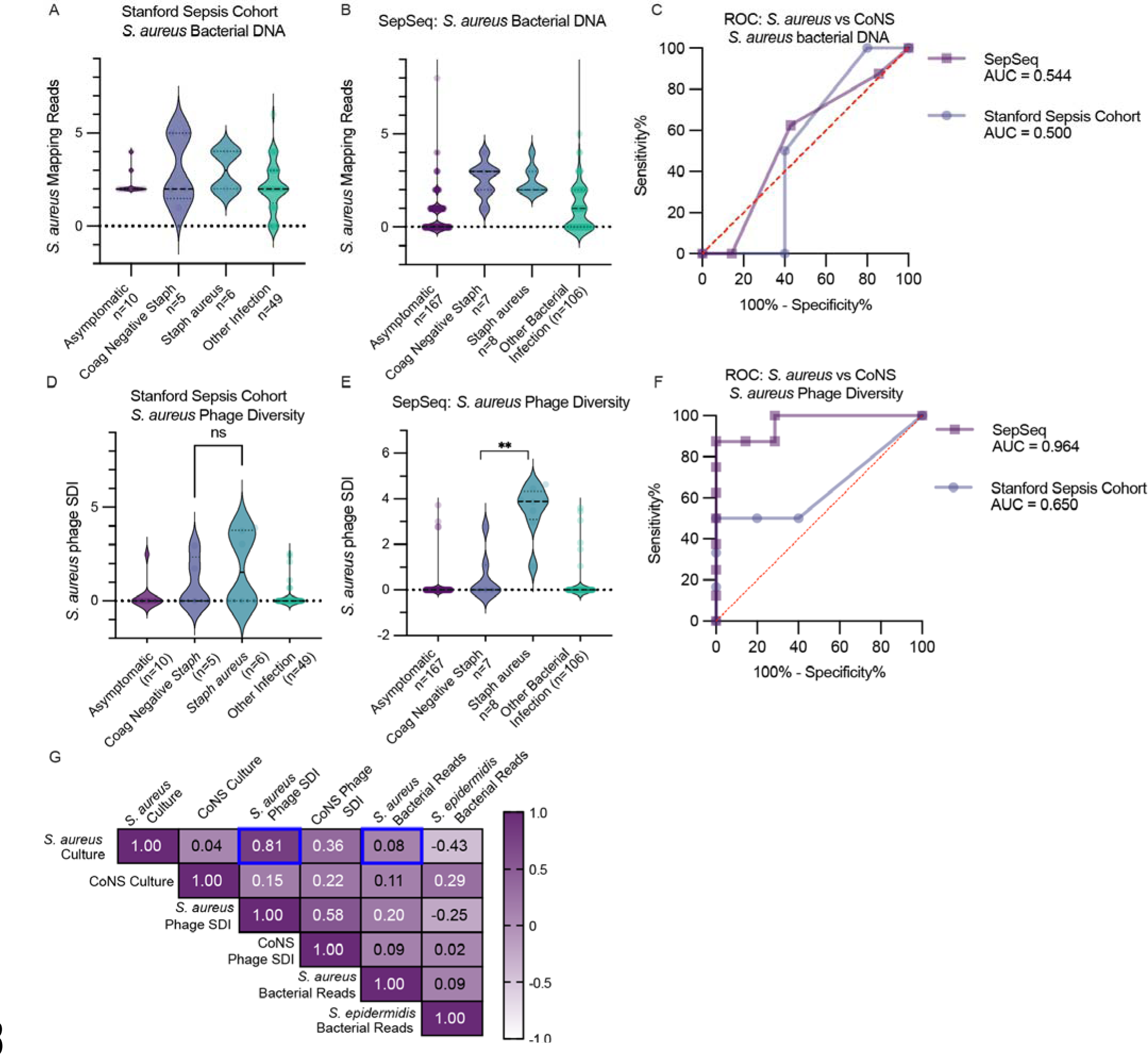
Phage can distinguish between Staphylococcus aureus and coagulase-negative Staphylococcus. A) Reads mapping to S. aureus bacterial genome in Stanford Sepsis Cohort by infection category (mean CoNS: 3, mean S. aureus: 3, Mann-Whitney test P > 0.99). One sample with both S. aureus and CoNS cultures excluded. B) Reads mapping to S. aureus bacterial genome in SepSeq by infection category (mean CoNS: 2.57, mean S. aureus: 2.38, Mann-Whitney test P = 0.844), C) ROC lot for S. aureus bacterial genome in distinguishing S. aureus culture vs CoNS culture (AUC Stanford Sepsis Cohort 0.500, 95% Confidence Interval 0.146 to 0.854, P > 0.99, AUC SepSeq 0.544, 95% Confidence Interval 0.225 to 0.858, P = 0.772), D) S. aureus phage SDI by infection category in Stanford Sepsis Cohort (mean CoNS: 0.78, mean S. aureus: 1.78, Mann-Whitney test P = 0.42) One sample with both S. aureus and CoNS cultures excluded, E) S. aureus SDI by infection category in SepSeq (mean CoNS: 0.553, mean S. aureus: 3.54, Mann-Whitney test P = 0.0012), F) ROC plot for S. aureus phage SDI in distinguishing S. aureus culture from CoNS culture (AUC Stanford Sepsis Cohort 0.667, 95% Confidence Interval 0.339 to 0.995, P = 0.337, AUC SepSeq 0.964, 95% Confidence Interval 0.877 to 1, P = 0.0026), G) Correlation matrix by culture status, phage diversity, and S. aureus mapping bacterial reads in SepSeq samples (S. aureus culture and S. aureus phage SDI Pearson R: 0.81, 95% Confidence Interval 0.736 to 0.861, P = 1.65e-029, CoNS Culture and S. aureus phage SDI Pearson R: 0.15, 95% Confidence Interval -0.031 to 0.32, P = 0.10, S. aureus culture and S. aureus Bacterial Reads Pearson R: 0.08, 95% Confidence Interval -0.098 to 0.254, P = 0.37, S. epidermidis Bacterial Reads and S. aureus culture Pearson R:-0.43, 95% Confidence Interval -0.759 to 0.060, P=0.08)

These data suggest that increased diversity of *S. aureus* phage is sensitive and specific for *S. aureus* infections. This relationship can also be seen in the correlation plot where *S. aureus* phage diversity correlates strongly with *S. aureus* infection, which was not the case for CoNS phage diversity and CoNS infection (Fig. 4G). Taken together, subgroups of phage, specifically *S. aureus* phage, are potential targets for comparing infections in related bacterial species where genetically similar pathogens complicate NGS analyses.

## Discussion

We report here that bacteriophage sequences are present in cfDNA from plasma and that this information can be used to noninvasively study bacteriophage ecology in bacterial infection. We demonstrate the utility of this approach in two independent data sets as well as in their associated asymptomatic controls, all generated using conventional NGS methods.

The data presented here indicate that all individuals have a circulating phageome. The phageome is reflective of bacteria associated skin and gut commensals - suggesting potential translocation of phage from the GI tract into circulation. However, low biomass of phage vs human DNA in cfDNA limit more specific analyses, such as generating quality phage contigs that could help to distinguish between integrated prophage and free phage DNA. The origins and functional significance of phage cfDNA in the human body await further investigation.

We found little global alteration in the phageome in bacterial sepsis. We demonstrate that phage can be used to study bacteriophage ecology in infection and to distinguish between closely related bacterial species that cause true infections and contamination. This potential was in the group of patients with *E. coli* sepsis, where we showed that phage ubiquitous across patients with *E. coli* sepsis tend to associate with toxin producing pathovariant hosts. We showed that bacteriophage may help recover lost resolution in plasma cfDNA when comparing closely related bacterial species such as *S. aureus* and CoNS. Future experimental studies in pre-clinical models are needed to inform our understanding of phage cfDNA in infection.

While the overrepresentation of phage corresponding to bacterial pathogens in sepsis appears robust (the parallel increase in bacterial cfDNA during sepsis is the foundation of various existing diagnostic platforms^11, 34, 45–47)^, its biological meaning is unclear. It is important to note that some phage may have broader bacterial host ranges than assumed^48^ – which may complicate analyses. Because bacteria may harbor multiple prophages^49^, our observed increase in diversity may reflect lysis by multiple phage due to rapid bacterial expansion or stress. However, other explanations are possible. Understanding phage dynamics over the course of an infection and in polymicrobial infections will increase robustness of phageome studies in infection. Further studies are needed to investigate how phage gain access to the bloodstream and how this varies by site of infection and bacterial burden. Larger studies to capture phage heterogeneity in the context of infection are needed before attempting to identify diagnostically useful phage targets.

This study has several limitations. One limitation is that both cohorts in this study were collected at the same location - the Stanford Emergency Department. However, sample preparation, year of acquisition, and sequencing methods differed between the groups. Additionally, because of the breadth of sepsis etiologies, each individual type of infection had relatively low numbers of samples, making comparisons challenging. Additionally, the asymptomatic control plasma samples – acquired commercially – were only from male donors, limiting the generalizability of the asymptomatic phageomes (Table S2). Another limitation is that low microbial cfDNA biomass limits such analyses to only known phages due to sparsity of reads. Though microbial cfDNA biomass may be higher at infection sites, we believe that the tradeoff of a noninvasive approach facilitates easier acquisition of patient samples and enables retrospective study in banked plasma samples from infected patients.

In summary, we showed here that there is a circulating phageome detectable in both asymptomatic and infected individuals using NGS sequencing of cfDNA, and that phage cfDNA can help to investigate bacterial and phage ecology in infection at the population level. We believe there is great promise in using phage to noninvasively study bacterial infection ecology, and that further exploration of bacteriophage ecology and dynamics in the context of bacterial infections will lay the foundation for future efforts to utilize phage in noninvasive diagnostics.

## Methods

### Sample collection

The Emergency Department Sepsis Biobank study protocol and informed consent (32851) was reviewed and approved by the University’s IRB (Stanford). Samples were collected from patients presenting to the Stanford University Hospital Emergency Department who triggered a sepsis alert, which included blood sample collection at the time of enrollment as well as collection of the results from standard of care microbiological testing they received during the first seven days of their admission. Patients were eligible to enroll if they triggered a sepsis alert, were 18 years of age or older, had a temperature of >38°C or <36°C and met at least one systemic inflammatory response syndrome criteria (heart rate >90 beats per minute; respiratory rate >20 breaths per minute or a partial pressure of carbon dioxide <32 mm Hg; white blood cell count of either > 12,000 cells μl^−1^ or <4,000 cells μl^−1^ and >10% bands).

Blood samples were collected from peripheral blood draw or indwelling venous catheter into BD Vacutainer EDTA blood collection tubes (Becton Dickinson, Franklin Lakes, NJ) at the same time as the patients were undergoing blood draw for standard of care blood cultures. In the vast majority of patients, this occurred prior to antibiotic administration. Samples were then stored at 4°C for no more than 72 hours, centrifuged at 1500 rpm for 10 min to obtain plasma, and stored thereafter at -80°C. Samples were deidentified using a unique code that could be linked to the patient identifier to later obtain gold standard microbiological testing results for each patient. Plasma samples from 61 patients with positive blood culture results were randomly selected for further phageome analyses as described below.

Asymptomatic control plasma was purchased from Innovative Research (IPLASK2E2ML Novi, Michigan).

### DNA extraction, library preparation, and NGS

DNA was extracted from 1 mL of each human plasma sample using the DNeasy Blood and Tissue kit (Qiagen, 69504, Hilden, Germany) according to the manufacturer protocol and recommendations. Extracted DNA was delivered to SeqMatic (Fremont, California) for DNA library preparation, which was performed using the Nextera XT library preparation kit (Illumina, FC-131-1096, San Diego, California) and then subsequent sequencing on the NovaSeq 6000 platform (1 x 100bp) at an average of 18.67 million reads per sample, which, due to low phage biomass, corresponds to a sequencing depth per phage of less than 1 for most phage detected.

Nuclease free water and phosphate buffered saline (PBS) were run through the DNeasy Blood and Tissue kit DNA extraction protocol and used as controls in the Nextera XT library preparation to characterize ambient DNA or contaminating DNA in kits used for DNA extraction or library preparation. After adapter trimming and human read removal (as described below), a blast search was done against the full nucleotide database. Blast output, including GI/subject sequence ID, Taxonomic ID, Name, Percent identical match, evalue, and query sequence, along with taxonomic category (Human, Eukaryotic, Bacterial, Viral, Bacteriophage, Plantaea, Other/Synthetic, and Undescribed Environmental) are provided in Supplemental File 4 (Negative Controls).xlsx.

The samples from which publicly available data were used (Karius Inc., accession PRJNA507824) were described in their original manuscript (DOI: 10.1038/s41564-018-0349-6) as being thawed, spiked with a synthetic normalization molecule controls, and centrifuged at 16,000 g for removal of human cells. The cfDNA in remaining plasma were extracted using the Mag-Bind cfDNA Kit in the Hamilton STAR liquid handling workstation. Libraries were created using the Ovation Ultralow System V2 kit and were sequenced on the Illumina NextSeq500 using a 75-cycle single end run at an average 24 million reads per sample. These samples were downloaded using the sratoolkit using the accession list from PRJNA507824 with the “—gzip” option.

Corresponding DNA template free library preparations were used by Karius Inc. in the preparation of their SepSeq manuscript – they found only samples by ID’s SRR8288617, SRR8288832, and SRR8288643 were in a batch with a single *Lactotoccus* phage contaminant – one asymptomatic control and two samples of different infection etiologies (*Staphylococcus* and *Streptococcus*).

### Sequencing read quality control

Raw sequencing read quality was assessed using the FASTQC software^35^. Trimming of adapter sequences and low quality individual reads was done using Trimmomatic 0.39^36^ with the following settings: SE -phred33 -threads 8 ILLUMINACLIP:TruSeq3-SE.fa:2:30:10:2:keepBothReads LEADING:3 TRAILING:3 SLIDINGWINDOW:4:15 MINLEN:36.

FASTQC reports were moved to a single directory and MultiQC^50^ was used on this directory with default options to assemble a quality report across all samples post-trimming to confirm average read quality above the default trimming threshold.

### Human read removal

Reads passing quality control were then mapped to human reference genome GRCh38 with Bowtie2 v2.4.4^51^ with default options generating an output sam file.

Reads that did not map were considered to be putative non-human reads. The output sam file was converted to a bam file using the samtools^52^ view -bS command. The non-mapping reads were extracted using samtools view with the options “-b -f 0×4”. The non-human reads were converted from a bam file to a FASTQ file using the samtools bam2fq command.

Seqtk was used to convert FASTQ files to FASTA files^53^ using the command “seqtk seq -A input.fastq > output.fasta”.

These sequences were then subjected to an additional layer of human read removal: BLAST v2.13.0^38, 39^ was then used to align putative non-human reads to all human-associated sequences (those with taxids 9606, 63221, or 741158) in NCBI Nucleotide (www.ncbi.nlm.nih.gov/nuccore), retrieved on 11/28/2021. The blast search was done with an output format of “6 qseqid sseqid pident length evalue stitle” with a culling limit of 1 and an e-value cutoff of 0.0005.

All reads with hits to human sequences with an e-value below 0.0005 were discarded and all remaining reads were considered to be non-human. This was done by using the command “awk ’{print $1}’ blast_output”.txt > toremove.txt” which extracted the sample read IDs (“qseqid”) from the blast output tables. Seqkit was then used to remove these reads and resulted in FASTA files which had been subjected to additional human sequence depletion, using the command “seqkit grep -v -- pattern-file toremove.txt sample.fasta > depleted_sample.fasta”.

### Superkingdom read annotation

Superkingdom distribution of non-human reads was done by nucleotide BLAST search against the full NCBI Nucleotide database (retrieved on 11/28/2021 as above). A negative taxid list containing primate taxids (9606, 63221, 741158) was provided to avoid any incorrect annotations that could result from residual human-origin reads not removed by the above rigorous human read depletion. A culling limit of 1 was used, and an evalue cutoff of 0.0005 was used. Output format was set by “6 qseqid sseqid length pident evalue staxids stitle”.

The distribution of superkingdoms was assessed by assessing proportions of all hit taxids which corresponded to eukaryotic, viral (mammalian or bacteriophage), and archaeal organisms. The lists of these taxids were obtained from the NCBI taxonomy database. The calculation of distribution was done in R Statistical Software (v4.1.2; R Core Team 2021^54^) with a summary table counting number of reads corresponding to each superkingdom created, and then corresponding proportions calculated for proportional visualization in Graphpad Prism.

### Bacteriophage annotations

The CPD was constructed from phage genomes retrieved from NCBI Nucleotide on 12/07/2021 (NCBI, US National Library of Medicine, retrieved as above); the search was restricted to viral sequences and used terms “phage” or “bacteriophage.” Duplicate sequences, or sequences of only ORFs, genes, proteins, regions, or modified genomes were excluded. This yielded 26,159 phage sequences, which were combined into a single FASTA file and used to generate a local phage BLAST database. Cleaned and processed reads were annotated by performing a BLAST alignment search of non-human reads against the CPD. The CPD BLAST database was used as the “-db”, wordsize was set to 28, an evalue of 0.0005 was set, and culling limit of 1. Output format was set to “6 qseqid sseqid pident length evalue stitle”. The reads with hits to phage sequences were then extracted from the FASTA file. This was done by using the command “awk ’{print $1}’ blast_output”.txt > tokeep.txt” which extracted the sample read IDs (“qseqid”) from the blast output tables.

Seqkit was then used to keep only these reads and resulted in FASTA files with only potential phage hits, using the command “seqkit grep --pattern-file tokeep.txt sample.fasta > subset_sample.fasta”.

All reads with bacteriophage hits below the significance threshold were then subjected to a BLAST alignment search limiting the subset of the NCBI Nucleotide database with human taxonomic IDs to identify bacteriophage matching reads with human homology. This was done using a BLAST database created from sequences with the taxid 9606 retrieved on 11/28/2021. The blast search was run with an evalue cutoff of 0.0005, culling limit of 1, 8 threads, and word size of 28. Output format was set using “6 qseqid sseqid pident length evalue stitle”. The reads with both phage hits and hits to human sequences with an evalue of 0.0005 or less were then removed from the FASTA files using command “awk ’{print $1}’ blast_output”.txt > toremove.txt” which extracted the sample read IDs (“qseqid”) from the blast output tables.

Seqkit was then used to remove these reads and resulted in FASTA files containing only sequences with hits to phage-only sequences with any sequences with human genome homology removed, using the command “seqkit grep -v --pattern-file toremove.txt subset_sample.fasta > depleted_subset_sample.fasta”.

The remaining sequences were subjected to a second BLAST search against the CPD with a word size of 28, use of 8 threads, an e-value cutoff of less than 0.0005, a culling limit of 10 to capture potential identities of various bacteriophages for reads covering highly conserved sequences across bacteriophages. Output included qseqid, sseqid, stitle, pident, length, and evalue.

The CPD, which includes phage and their bacterial host(s), was created using annotations from virus-host DB^40^, from the phage NCBI taxonomy entry, by phage name, and manual curation from manuscripts associated with the phage sequence submission. Phage with only genus level host detail available were noted as such. Further characterization of host bacterial qualities was performed through literature search. Accessions from the Cenote Human Virome Database^32^ (CHVD) gut phages were used to identify proportions of annotated, uncharacterized phage that have been found to be gut phages. The CPD is publicly available at https://doi.org/10.5281/zenodo.7154236. Of note, the CPD is reflective of a bias in nucleotide database enrichment for human associated pathogens and their phages, and as such underrepresents environmental phages.

### Interpretation of phage annotations

Phage sequence hits were analyzed using R. Phage accession IDs were extracted and the taxize package was used to assign corresponding taxonomic lineage information^55^. Phage diversity was calculated using the SDI ^41^ through the R package vegan^56^. R code for the use of the CPD along with blast outputs for interpretation of phageome representation of phage families and host genus/characteristics is available at (https://doi.org/10.5281/zenodo.7734114).

Pearson dissimilarity matrix was generated using the R package factoextra^57^. Genetically defined taxonomic lineages were used from a publicly available database^43^ for alternative analyses. The subset of E. coli phages was analyzed with the genetically defined lineages from INPHARED database^43^ used in place of the phage family identities present in the CPD for creating family summary tables and subsequent comparisons in GraphPad Prism. Sankey diagram of phage reclassification of identified E. coli phages was created using SankeyMATIC, which show on the left the original NCBI taxonomy defined taxonomic family, and on the right the taxonomic family identified in the INPHARED database.

### Bacterial mapping

Bacterial reads were assessed through mapping to the reference *S. aureus* bacterial genome NC_007795.1 using Bowtie2 v2.4.4^51^ with default settings. Number of mapping reads were obtained using the samtools view command, with flag -F 0×04 used to obtain a coverage output from the mapped SAM file output.

### Statistics

Statistics were performed using Prism 9.3.1 (GraphPad Software, San Diego, California). This included Mann-Whitney tests, Kruskal-Wallis test with Dunn’s multiple comparisons, two way ANOVA with multiple comparisons, Brown-Forsythe and Welch Anova Test with multiple comparison, unpaired t-test, receiver-operator curve generation, and Pearson correlations. Please see figure legends for specific statistical tests performed in each figure.

## Materials & Correspondence

All commercially obtained materials used in this study have catalog numbers noted in relevant methods sections. The entire volume of human plasma samples were used as described in the methods and therefore cannot be made available. Any correspondence addressed to Dr. Paul L Bollyky at.

## Supporting information

Supplemental File 1 (phage dictionary)

Supplemental File 3 (Metadata)

Supplemental File 2 (coliphage characteristics)

Supplemental File 4 (Negative controls)

## Acknowledgements

We would like to thank Tim Blauwkamp, Sivan Bercovici, and Nicholas Noll (Karius Inc) for their assistance providing additional metadata for the SepSeq dataset. PLB is supported by the NIH (R01 HL148184-01, R01 AI12492093, R01 DC019965), the Cystic Fibrosis Foundation, and a grant from the Emerson Collective. NLH is supported by the NSF GRFP. LJB is supported by the NIH (T32 HL129970-06). SY is supported by the NIH (R01 AI153133, R01 AI137272, R01 AI138978)

## Author Contributions

N.L.H., L.J.B., N.R.M., G.K., S.Y., and P.L.B. designed the study. N.L.H., L.J.B., and N.R.M performed experiments. N.L.H., L.J.B., G.K., N.R.M., S.Y., A.S.B., C.Y.C. and P.L.B. analyzed data. N.L.H., L.J.B., and P.L.B. wrote the manuscript.

## Data Availability

Sequencing data with human reads removed have been deposited into NCBI SRA under bioproject PRJNA860730.

Publicly available data utilized: The SepSeq study data has been previously published under bioproject PRJNA507824. No new computational tools were developed as part of this study.

Infection etiology metadata associated with samples sequenced for this study are included in Supplemental File 3 (Metadata).xlsx.

The Curated Phage Database (CPD) FASTA file used for creating the Blast database is publicly available at: https://doi.org/10.5281/zenodo.7154236.

All associated supplementary files have additionally been made publicly available at: https://doi.org/10.5281/zenodo.7644125.

## Code Availability

The R code used to summarize BLAST phageome annotations with the CPD have been made publicly available at (https://doi.org/10.5281/zenodo.7734114). These include an R markdown file detailing processing of BLAST outputs to create phage hit tables across all samples, and subsequent use of the CPD to summarize representation of phage taxonomic families and known bacterial host characteristics. A phage hit table for our sequenced samples are available along with this R code and can be used to recreate phage summary tables as well as for calculation of diversity using the R package “vegan”. Processing of raw data, removal of human reads, and BLAST annotations were done using existing software and are described in the relevant methods sections.

## Figures

**Figure S1.**
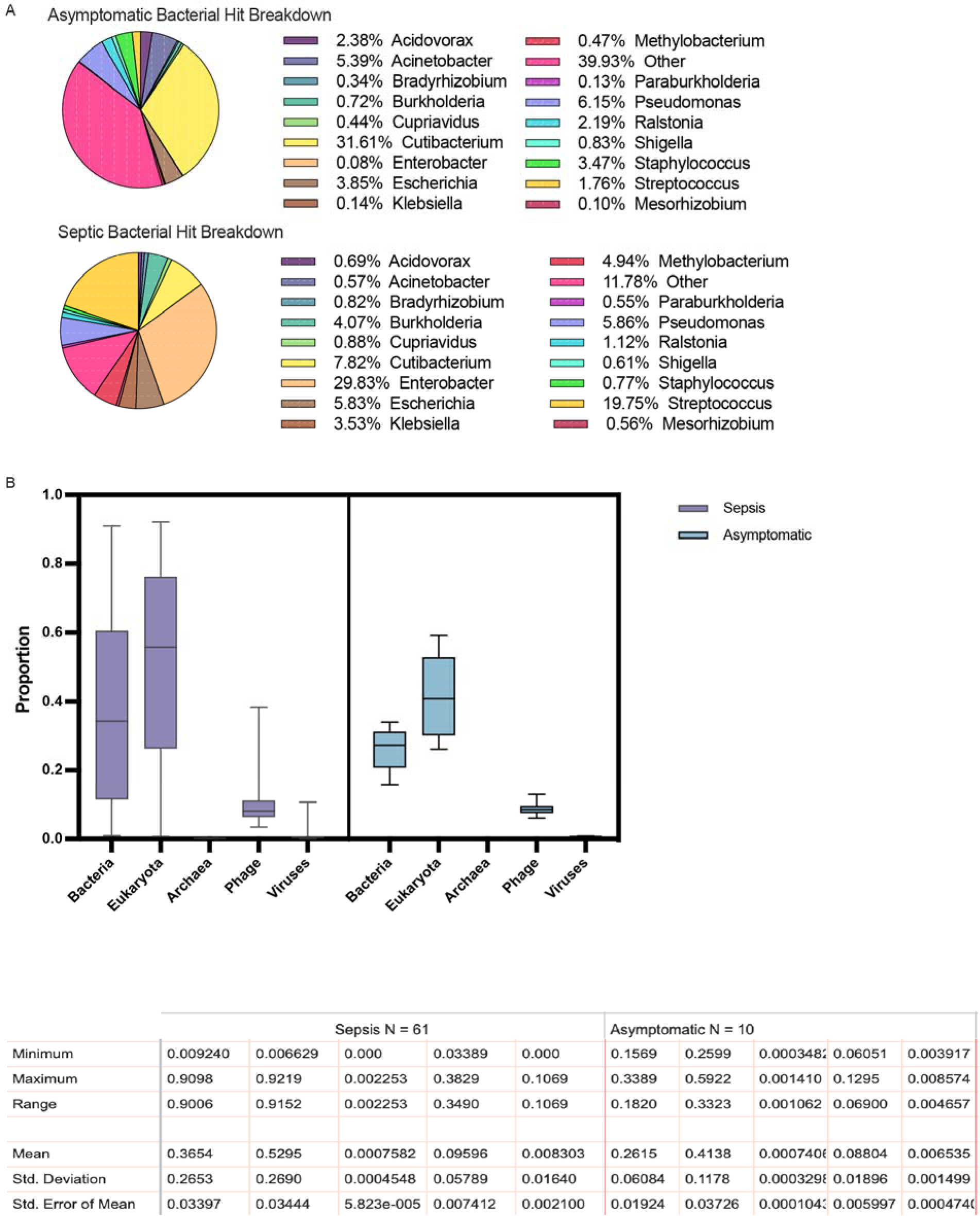
Non-Human reads in Asymptomatic and Septic individuals. A)Average proportion of bacterial hit genus in asymptomatic and septic nonhuman cfDNA as identified by BLAST search. B) Bar plots of proportions of non-human read identities by BLAST search with a table of descriptive statistics for samples from both Septic and Asymptomatic individuals.

**Figure S2.**
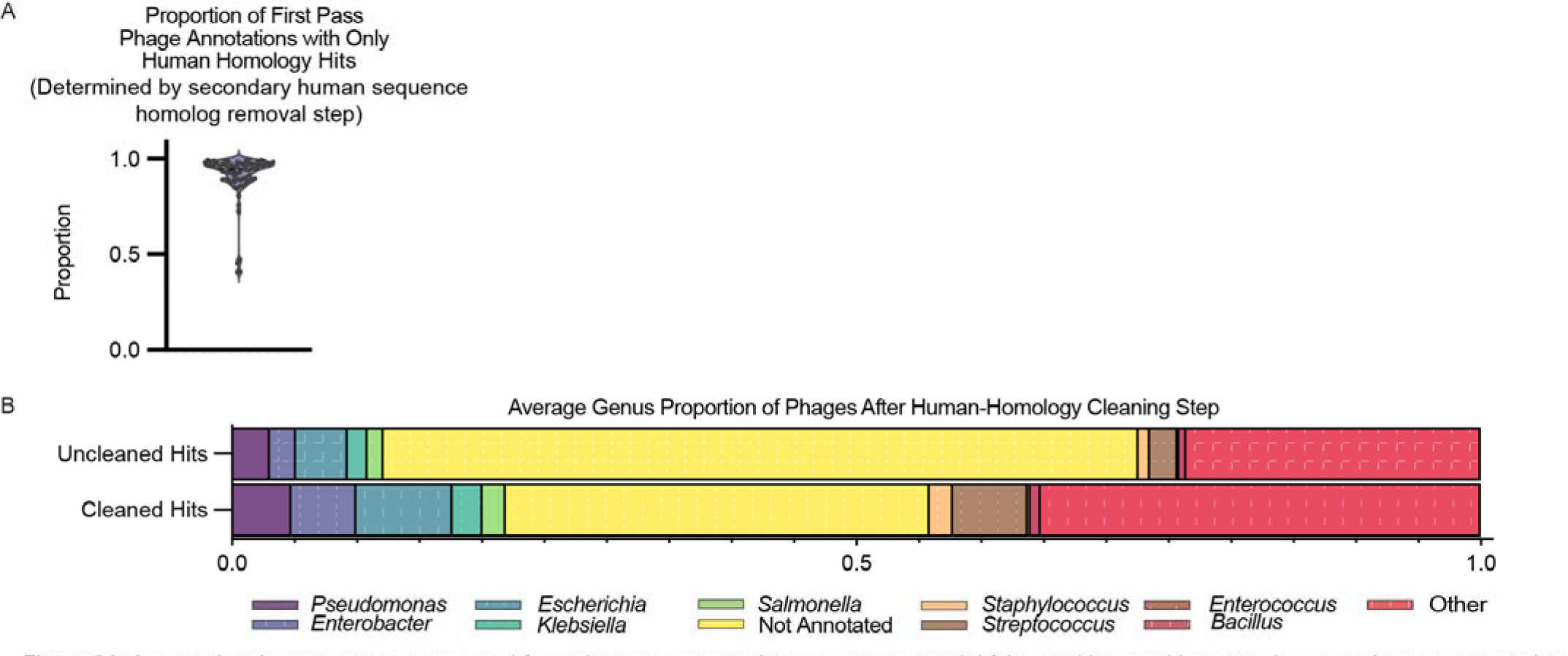
A secondary human sequence removal from phageome annotations removes potential false positives and impacts phageome host representation. A) Proportions of bacteriophage hits removed in secondary human sequence homolog removal step mean 0.907 SD 0.128, B) Average distribution of unique phages by bacterial host genus with and without secondary human sequence homology removal. Uncleaned Hits, mean proportions and SD *(Pseudomonas:* mean 0.029 SD 0.30, *Enterobacter* mean 0.021 SD 0.015, *Escherichia* mean 0.042 SD 0.038, *Klebsiella* mean 0.015 SD 0.020, *Salmonella* mean 0.013 SD 0.009, Not Annotated mean 0.603 SD 0.070, *Staphylococcus* mean 0.009 SD 0.011, *Streptococcus* mean 0.014 SD 0.036, *Enterococcus* mean 0.001 SD 0.002, *Bacillus* mean 0.006 SD 0.007, Other mean 0.245 SD 0.052), Cleaned Hits, mean proportions and SD *(Pseudomonas:* mean 0.047 SD 0.058, *Enterobacter mean* 0.050 SD 0.058, *Escherichia* mean 0.076 SD 0.065, *Klebsiella* mean 0.023 SD 0.036, *Salmonella* mean 0.014 SD 0.016, Not Annotated mean 0.35 SD 0.100, *Staphylococcus* mean 0.020 SD 0.040, *Streptococcus* mean 0.032 SD 0.076, *Enterococcus* mean 0.002 SD 0.004, *Bacillus* mean 0.012 SD 0.015, Other mean 0.373 SD 0.123)

**Figure S3.**
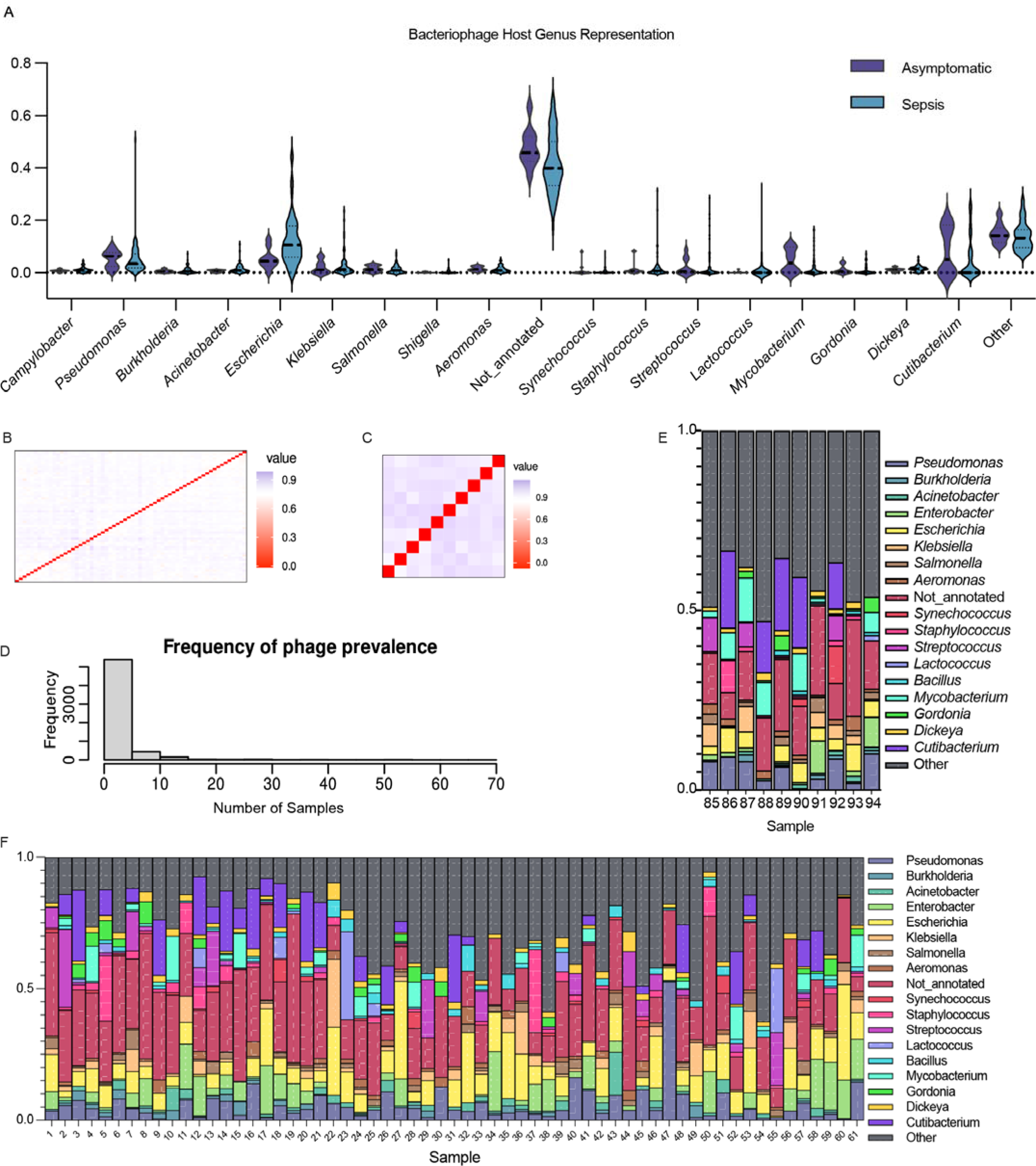
Phage bacterial host distribution does not change in sepsis, though individual variation remains. A) Violin plot of bacteriophage host genus proportions between Asymptomatic and Septic patient samples, associated statistics are in Table S1. B) Heatmap of Pearson dissimilarity matrix between patients with sepsis. C) Heatmap of Pearson dissimilarity matrix between asymptomatic controls. D) Histogram of prevalence across sequenced samples of each phage. E) Phage bacterial host genus proportions per asymptomatic patient. F) Phage bacterial host genus proportions per septic patient.

**Figure S4.**
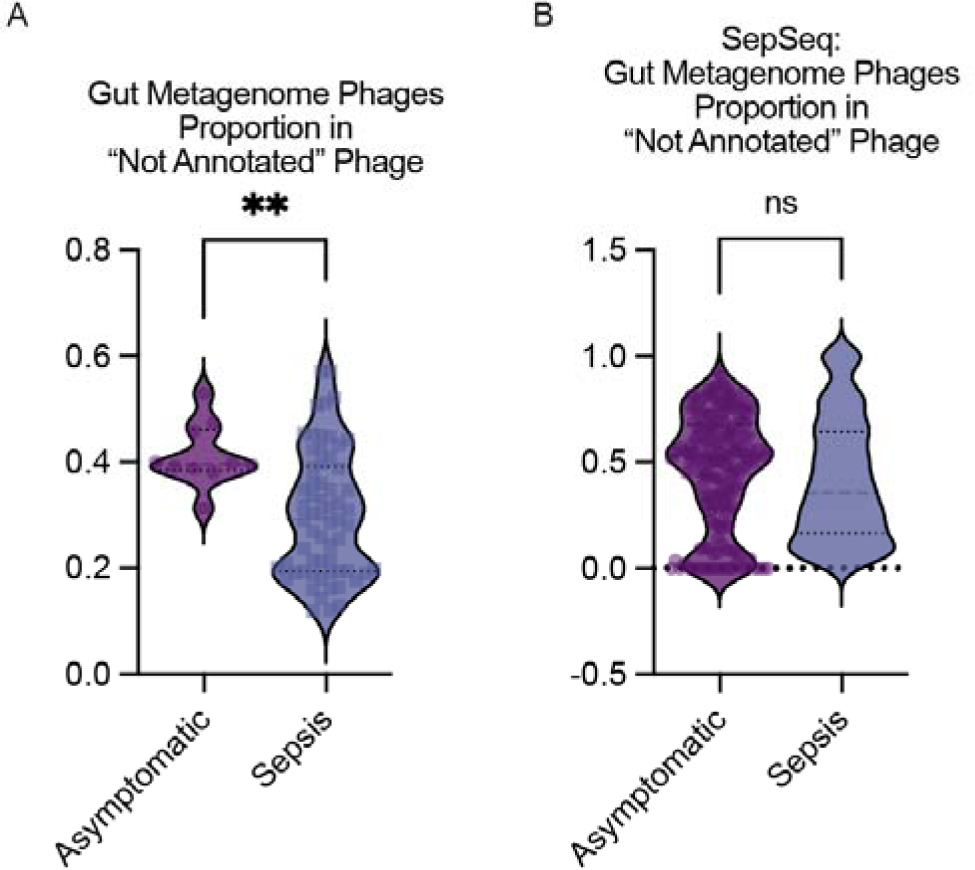
Proportion of “Not Annotated” Phages from CHVD Gut Metagenome Phages. A) Proportion of “Not Annotated” Phages from Gut Metagenome Phages in Stanford Sepsis Cohort (Asymptomatic mean: 0.413 SD: 0.060, Sepsis mean: 0.300 SD:0.119. Unpaired t test, P = 0.0046) B) Proportion of “Not Annotated” Phages from Gut Metagenome Phages in SepSeq cohort (Asymptomatic mean: 0.464 SD:0.271, Sepsis mean: 0.418, SD:0.29. Unpaired t test, P = 0.101)

**Figure S5.**
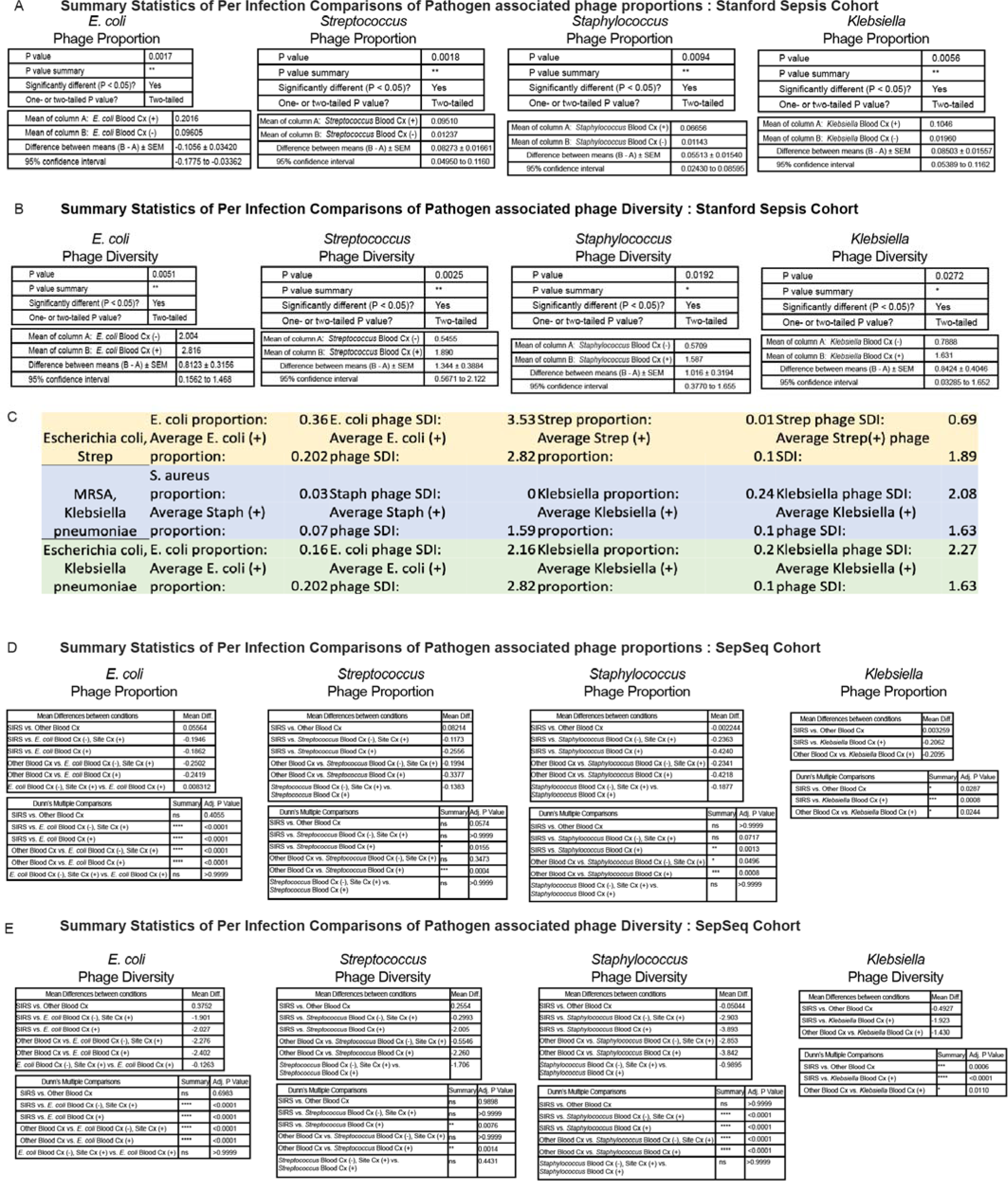
Statistical summaries showing pathogen specific phage proportion is higher in samples from patients with those infections. A) In Stanford Sepsis cohort, Proportion of infection specific phage in samples with a positive culture for that pathogen vs samples with a different identified pathogen, Statistics shown for Mann-Whitney tests between groups, including mean difference of means, 95% Cl, and p value. B) In Stanford Sepsis cohort, Diversity of infection specific phage in samples with a positive culture for that pathogen vs samples with a different identified pathogen. Statistics shown for Mann-Whitney tests between groups, including mean difference of means, 95% Cl, and p value C) In Stanford Sepsis cohort, comparison of proportion and phage SDI of three polymicrobial infections to the averages for those infection etiologies. D) In SepSeq cohort, Proportion of infection specific phage in samples with positive blood culture, positive site of infection culture only, other identified pathogen, or SIRS. Statistics shown for Kru ska I-Wallis test Dunn Multiple Comparisons between groups, including mean difference of means, 95% Cl, and p value **E)** In SepSeq cohort, Diversity of infection specific phage in samples with positive blood culture, positive site of infection culture only, other identified pathogen, or SIRS. Statistics shown for Kruskal-Wallis test Dunn Multiple Comparisons between groups, Including mean difference of means, 95% Cl, and p value

**Figure S6.**
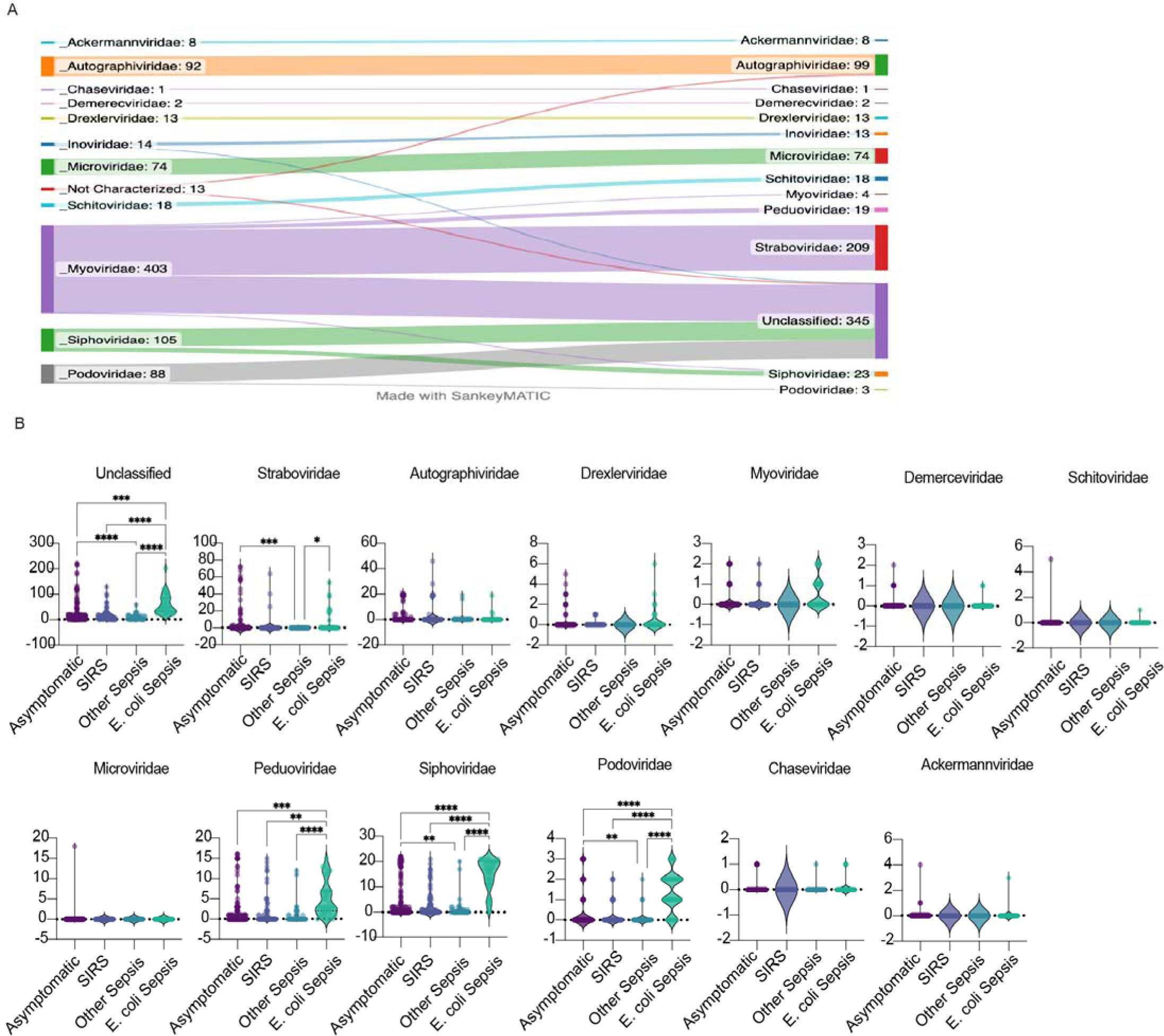
Number of E. coli phages by genetically classified taxonomic phage family. A) Sankey diagram of taxonomic family classifications from NCBI Taxonomy classification (Left) to genetically classified family (Right) B) Number of E. coli phages by genetically classified taxonomic phage family tested by Brown-Forsythe and Welch Anova Test with multiple comparisons.

**Figure S7.**
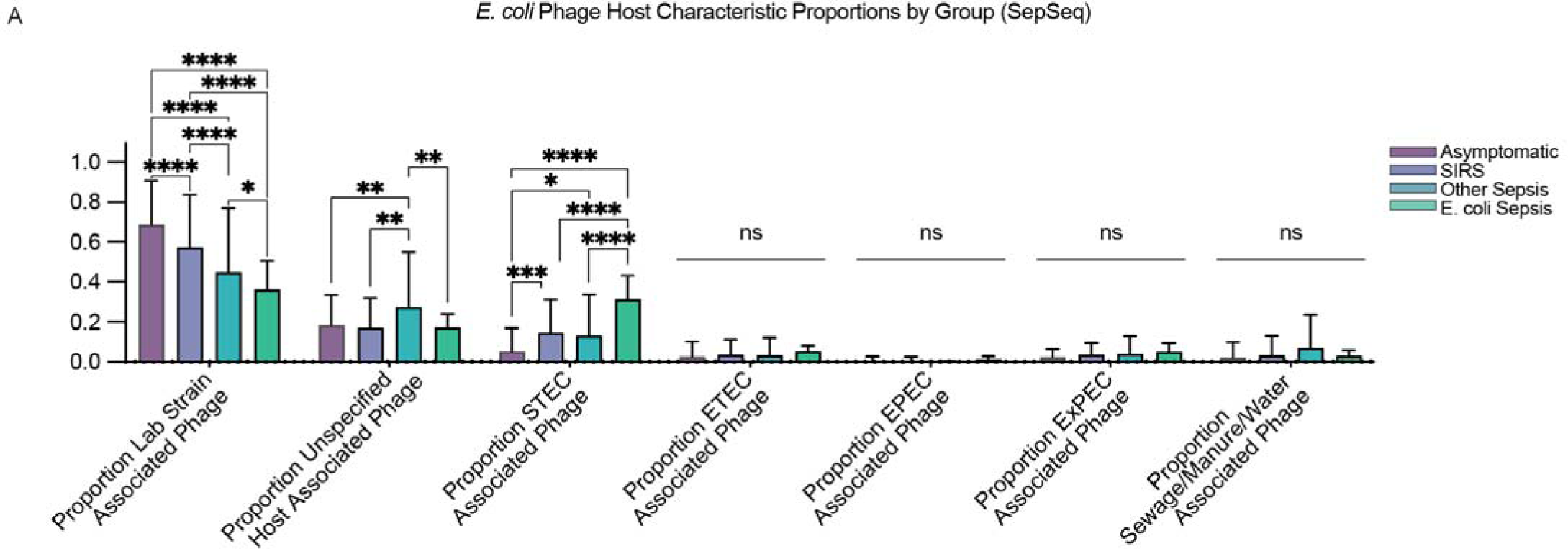
*E, coli* Phage Host Characteristic Proportions. Differences in proportion of *E. coli* phage host characteristics from the literature are shown. Analyzed by two way ANOVA with multiple comparisons. Shown on plots are ns if no differences were found across characteristic proportion or between conditions.

**Table S1:** Mann-Whitney test summary of Phage SDI between Asymptomatic and Septic patient samples. All genus specific SDIs had an adjusted p value greater than 0.05.

|  | Discovery? | P value | Mean rank of Asymptomatic | Mean rank of Sepsis | Mean rank diff. | Mann-Whitney U | Adj. p value |
| --- | --- | --- | --- | --- | --- | --- | --- |
| Campylobacter | No | 0.113756 | 26.75 | 38.07 | -11.32 | 212.5 | 0.57447 |
| Pseudomonas | No | 0.497073 | 40.75 | 35.81 | 4.935 | 267.5 | 0.772375 |
| Burkholderia | No | 0.775651 | 34.75 | 36.78 | -2.032 | 292.5 | 0.870453 |
| Acinetobacter | No | 0.175584 | 28.1 | 37.85 | -9.755 | 226 | 0.591131 |
| Escherichia | No | 0.003646 | 19.15 | 39.3 | -20.15 | 136.5 | 0.073656 |
| Klebsiella | No | 0.679095 | 33.9 | 36.92 | -3.019 | 284 | 0.857357 |
| Salmonella | No | 0.560555 | 40.1 | 35.92 | 4.181 | 274 | 0.808801 |
| Shigella | No | 0.421434 | 32.1 | 37.21 | -5.11 | 266 | 0.709415 |
| Aeromonas | No | 0.214896 | 44.2 | 35.26 | 8.942 | 233 | 0.620128 |
| Not annotated | No | 0.044566 | 48.8 | 34.52 | 14.28 | 187 | 0.300079 |
| Synechococcus | No | 0.844428 | 35.35 | 36.69 | -1.335 | 298.5 | 0.872229 |
| Staphylococcus | No | 0.863593 | 35.4 | 36.68 | -1.277 | 299 | 0.872229 |
| Streptococcus | No | 0.654721 | 39.1 | 36.08 | 3.019 | 284 | 0.857357 |
| Lactococcus | No | 0.774446 | 35.3 | 36.69 | -1.394 | 298 | 0.870453 |
| Mycobacterium | No | 0.030548 | 48.9 | 34.5 | 14.4 | 186 | 0.300079 |
| Gordonia | No | 0.409273 | 41.1 | 35.76 | 5.342 | 264 | 0.709415 |
| Dickeya | No | 0.143066 | 27.45 | 37.96 | -10.51 | 219.5 | 0.577986 |
| Cutibacterium | No | 0.301805 | 42.2 | 35.58 | 6.619 | 253 | 0.709415 |
| Other | No | 0.336353 | 42.5 | 35.53 | 6.968 | 250 | 0.709415 |

**Table S2:**
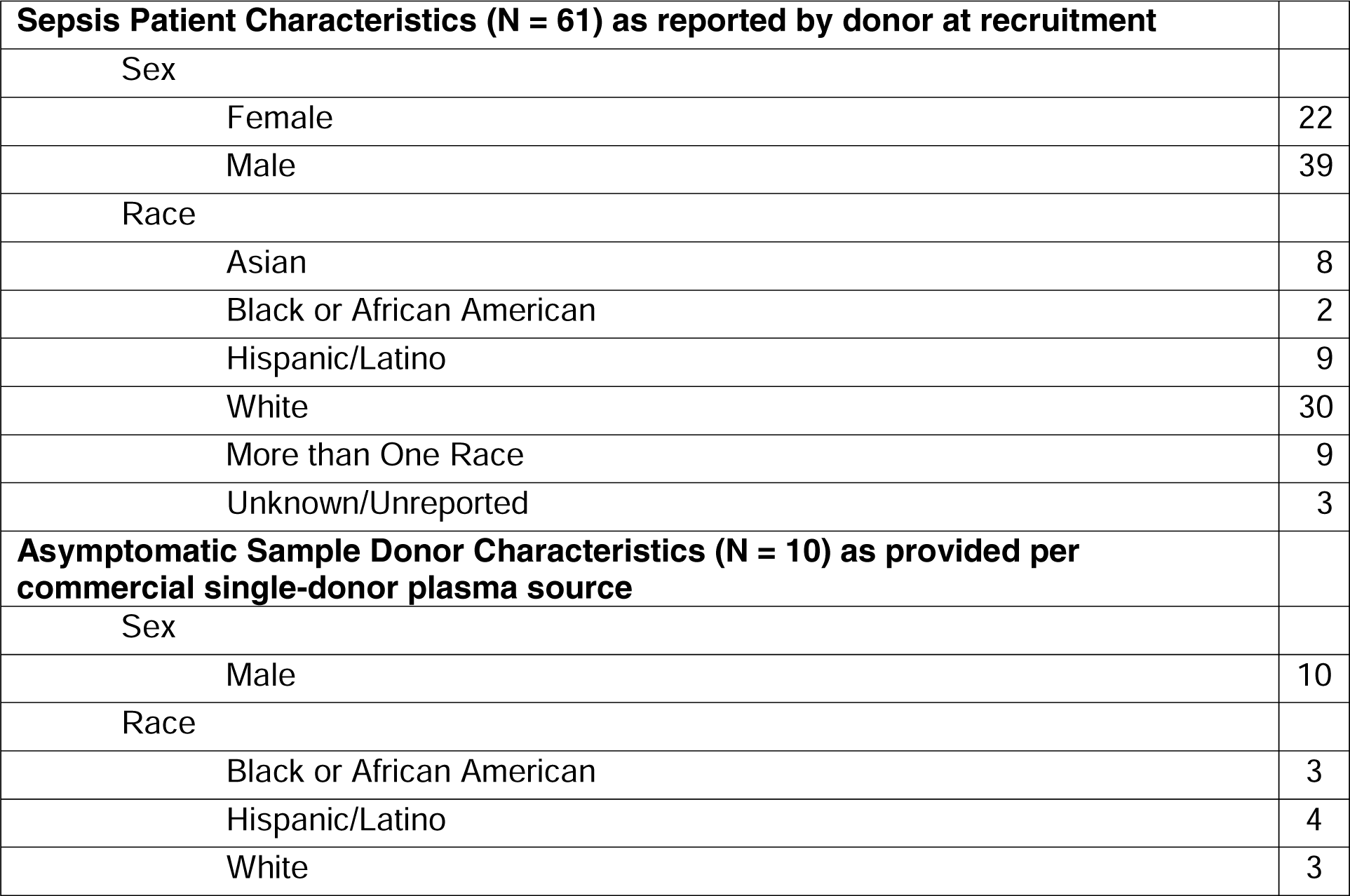
Plasma Sample Donor Demographic Characteristics

